# A synthetic non-histone substrate provides insight into substrate targeting by the Gcn5 HAT and sirtuin HDACs

**DOI:** 10.1101/345637

**Authors:** Anthony Rössl, Mong-Shang Lin, Michael Downey

## Abstract

Gcn5 and sirtuins are highly conserved HAT and HDAC enzymes that were first characterised as regulators of gene expression. Although histone tails are important substrates of these enzymes, these proteins also target many non-histone substrates that participate in diverse biological processes. The mechanisms used by these enzymes to choose their non-histone substrates is unclear. In this work, we use a unique synthetic biology approach in *S. cerevisiae* to demonstrate that a shared target sequence can act as a determinant of substrate selection for Gcn5 and sirtuins. We also exploit this system to define specific subunits of the Gcn5-containing ADA complex as regulators of non-histone acetylations proteome-wide.

## INTRODUCTION

The Gcn5 histone acetyltransferase (HAT) is a member of the GNAT family of acetyltransferase enzymes. It functions in the context of a highly conserved protein complex called SAGA that contains at least 19 unique subunits. These subunits can be grouped into functional submodules that together regulate important aspects of eukaryotic gene transcription (Lee et al. 2011; Spedale et al. 2012; Han et al. 2014; Helmlinger and Tora 2017). Besides Gcn5, the HAT submodule contains Ada2, Ada3 and Sgf29 (Lee et al. 2011). The proteins of the HAT submodule also function in a distinct complex termed ADA that includes Ahc1 and Ahc2 (Eberharter et al. 1999; Lee et al. 2011). SAGA deubiquitylation (DUB) and TATA-binding protein (TBP) regulatory modules (consisting of Spt3 and Spt8) mediate deubiquitylation of H2B K123 and recruitment of the TBP to gene promoters, respectively (Warfield et al. 2004; Ingvarsdottir et al. 2005; Lee et al. 2005; Sermwittayawong and Tan 2006; Laprade et al. 2007). A core structural model that includes subunits shared with general transcription factor TFIID serves as a scaffold for the SAGA complex (Lee et al. 2011; Han et al. 2014).

Ada2 and Ada3 play important roles in promoting Gcn5 activity towards histone substrates, particularly in the context of nucleosomes (Marcus et al. 1994; Balasubramanian et al. 2002; Sterner et al. 2002b). While Sgf29 is largely dispensable for Gcn5 activity *in vitro*, it plays a critical role in global histone acetylation *in vivo*, as *sgf29*Δ cells show decreased acetylation of histone H3 K9, K14 and K18, paralleling what is observed for *gcn5*Δ and *ada3*Δ mutants (Bian et al. 2011). This function of Sgf29 is likely due to the ability of its Tudor domain to bind to methylated H3 K4 (Bian et al. 2011). SAGA integrity is also required for histone acetylation *in vivo*, as deletion of genes encoding scaffold elements Spt20 or Spt7 results in decreased global H3 acetylation (Peng et al. 2008).

Like HATs, histone deacetylase (HDAC) enzymes are grouped into families based on common structural and biochemical characteristics. The NAD+ dependent family of sirtuin HDACs, consisting of Sir2 and Hst1-Hst4, are conserved enzymes that can be inhibited with a byproduct of their reactions called nicotinamide (Brachmann et al. 1995; Wierman and Smith 2014). Sirtuins Hst3 and Hst4 deacetylate H3 K56 (Celic et al. 2006; Maas et al. 2006), which is important for DNA repair and the maintenance of genome integrity (Celic et al. 2008; Che et al. 2015). Hst4 also localizes to the mitochondria where it regulates protein deacetylation in response to biotin starvation (Madsen et al. 2015). Sir2 and Hst1 function in gene silencing and transcriptional control at select genomic loci (Imai et al. 2000; Landry et al. 2000; Mead et al. 2007; Froyd and Rusche 2011). Finally, Hst2 is the only cytoplasmic sirtuin (Perrod et al. 2001; Wilson et al. 2006) and its function remains poorly characterized.

Although acetylation was originally characterized as a histone modification and regulator of gene transcription, thousands of non-histone substrates have been described using high-throughput approaches in organisms from bacteria to humans (Choudhary et al. 2009; Weinert et al. 2011; Henriksen et al. 2012; Downey et al. 2015). In yeast, at least one-third of all proteins are acetylated (Downey and Baetz 2016). While regulation of histone acetylation and deacetylation activities is mediated by temporal and spatial changes in HAT and HDAC recruitment to specific chromatin loci, the factors governing selection of non-histone substrates are less clear.

In previous work, we used SILAC labeling of yeast cells coupled with affinity enrichment of acetylated peptides and mass spectrometry to uncover candidate substrates of the Gcn5 and Esa1 HATs and the sirtuin family of HDACs (Downey et al. 2015). Analysis of high-confidence candidate targets uncovered preferred amino acid motifs surrounding regulated acetylated lysines (Downey et al. 2015). Intriguingly, there were similarities between these “consensus” target sequences for Gcn5 and sirtuin enzymes, with S-x-K(ac)-K/R-P being preferred for both enzymes. This shared sequence was distinct from that previously identified for Gcn5 (Rojas et al. 1999), and from that of Esa1, which bore significant resemblance to the glycine-rich H4 tail (Downey et al. 2015).

Here, we used a synthetic biology approach to demonstrate that this shared sequence is sufficient to confer Gcn5- and sirtuin-regulated acetylation *in vivo.* A fusion protein containing GFP fused to variants of the consensus sequence, in conjunction with an antibody directed against that acetylated consensus, serve as a toolkit to probe sirtuin and Gcn5 function *in vivo.* Our work with this toolkit points to a model where Gcn5 activity towards lysine residues within preferred sequence contexts depends on association with Ada2 and Ada3 but is largely independent of other SAGA proteins.

## RESULTS AND DISCUSSION

### A shared consensus sequence predicts Spt2 as a novel target of Gcn5 and sirtuins

We previously identified a shared consensus sequence of S-x-K(ac)-K/R-P for Gcn5 and sirtuin enzymes by carrying out SILAC-based acetylome analyses for *gcn5*Δ and *hst1*Δ *hst2*Δ *sir2*Δ triple mutant cells. We first wondered if this sequence could be used to predict new sites regulated by these enzymes. We focused on the S-S-K(ac)-R-P sequence, which represents the most frequently observed amino acids surrounding Gcn5-depdendent acetylations, corrected for relative amino acid frequencies in yeast. Four proteins contain an exact match: Spt2, Far10, Afr1 and Ydr249c **(Fig. 1A)**. We were able to generate GFP-tagged versions of Spt2, Far10 and Ydr249c. Spt2 is a member of transcriptional regulator that physically interacts with the SWISNF chromatin remodelling complex (Perez-Martin and Johnson 1998). Far10 is a member of the conserved STRIPAK complex that mediates pheromone and TORC2-dependent signaling pathways in yeast (Bloemendal et al. 2012; Pracheil and Liu 2013). Ydr249c is a largely uncharacterized protein (Cherry et al. 2012). We immunopurified these GFP-fusion proteins and tested for reactivity with monoclonal antibodies recognizing acetylated lysine in the context of defining features of the S-x-K(ac)-K/R-P sequence (See Materials & Methods). In this experiment, *FAR10*-GFP and *YDR249C*-GFP were expressed from the inducible *GAL* promoter (Longtine et al. 1998) to allow recovery of a sufficient level of protein, whereas *SPT2*-GFP was expressed at sufficient levels under its endogenous promoter. We observed no evidence of Far10-GFP and Ydr249c-GFP acetylation **(Supplemental Fig. S1)**. In contrast, Spt2-GFP showed reactivity with monoclonal anti-acetyllysine antibody following recovery from cells treated with sirtuin inhibitor nicotinamide **(Fig. 1B**), and expression of GFP-tagged Spt2 mutated for the lysine residue (K166) within its S-S-K-R-P consensus sequence completely eliminated the signal **(Fig. 1B,C)**. Finally, as predicted from the consensus sequence, acetylation was dependent on *GCN5* **(Fig. 1D)**. Altogether, these data are consistent with Gcn5-regulated acetylation of Spt2-GFP K166 and the reversal of this modification by sirtuin enzymes. The data highlight that acetylation consensus sequences derived from high-throughput mass spectrometry data can be used to identify novel targets for HAT and HDAC enzymes. Notably, in contrast to our sequence-specific monoclonal antibodies, a commonly used pan-acetyllysine antibody (Cell Signaling 9441) did not detect regulated acetylations on Spt2-GFP (data not shown). The monoclonal antibodies described in this work (see Materials and Methods) could function as a new tool to study Gcn5 and sirtuin-regulated acetylations in yeast.

**Figure 1:**
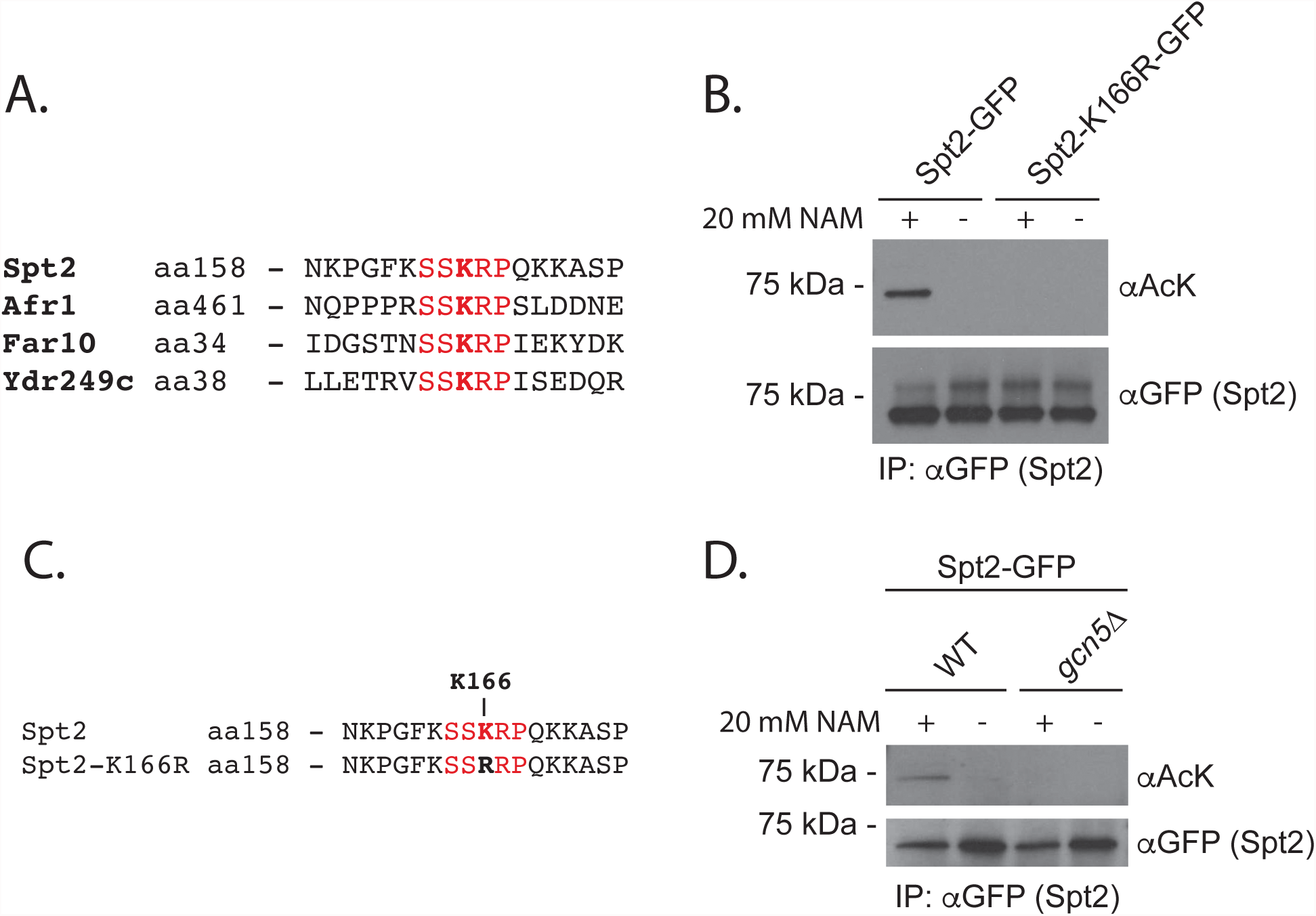
Consensus target sequences predict Spt2 as a candidate Gcn5 and sirtuin target. **A)** Alignment of yeast proteins containing an exact match to the SSKRP consensus sequence. **B)** WT or Spt2-GFP K166R was immunoprecipitated in the presence or absence of nicotinamide treatment (20 mM, 2 generations). Eluates were separated via SDS-PAGE, transferred to PVDF membrane and probed with aGFP or monoclonal antibody developed to recognize the acetylated SSKRP sequence**. C)** Alignment of Spt2 target sequences showing the K-R mutation used in (B). **D)** Acetylation of Spt2-GFP was measured in a *gcn5*Δ strain in the presence and absence of nicotinamide.

Of Spt2, Far10 and Ydr249c, only Spt2 showed regulated acetylation of its S-S-K-R-P consensus sequence via Gcn5 and sirtuins **(Fig 1B, Supplemental Fig. S1)**. As such, presence of the consensus sequence alone is not sufficient to confer acetylation. Of the three candidate targets, only Spt2 has demonstrated localization to the nucleus (Kruger and Herskowitz 1991). Thus, it is possible that nuclear localization is required for acetylation by Gcn5.

### A synthetic non-histone substrate is acetylated in vivo

In order to further probe the contribution of the shared Gcn5/sirtuin sequence to protein acetylation, we asked whether the addition of this sequence to a non-substrate would be sufficient to confer acetylation *in vivo*. We fused increasing numbers of S-S-K-R-P consensus sequence to GFP **(0X-3X; Fig. 2A)**. We chose GFP because it does not react with anti-acetyllysine antibodies in IP-Western experiments (see below) and can localize throughout the cell (Niedenthal et al. 1996; Yen et al. 2001). We expressed these fusion constructs or GFP alone from a constitutive ADH1 promoter and used an IP-Western strategy to recover and compare their acetylation using our S-S-K(ac)R-P-reactive monoclonal antibodies. We detected acetylation on our fusion constructs, but not GFP alone **(Fig. 2B).** Moreover, the acetylation signal increased with the number of consensus repeats (**Fig. 2B, Supplemental Fig. S2A)**.

**Figure 2:**
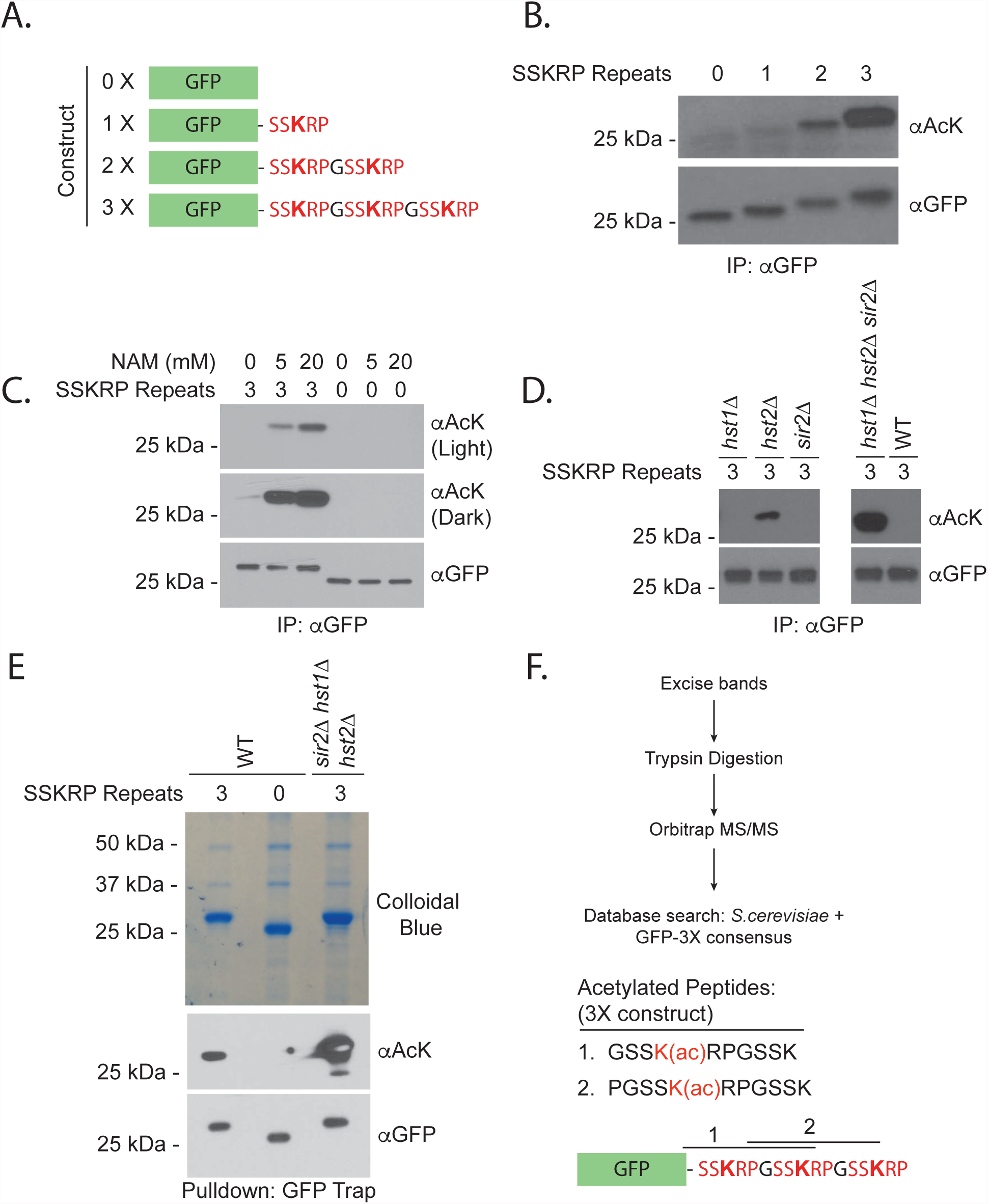
A synthetic acetylation substrate is acetylated *in vivo*. **A)** Synthetic fusion constructs used for acetylation analyses. **B)** GFP fusions were purified from strains expressing the constructs in (A) using an antibody against GFP. Eluates were analyzed via SDS-PAGE and probed either with αAcetyllysine or αGFP following Western blotting. Diluted forms (1/25) of immunoprecipitated protein samples were loaded for GFP detection. **C)** The indicated constructs were purified from strains treated with 0, 5 or 20 mM nicotinamide for 30 minutes and analyzed as in (B). **D)** The indicated constructs were purified from the sirtuin mutant strains shown and analyzed as in (B). Single and triple mutant blots were cropped from the same image. **E)** Indicated constructs were purified from the strains shown using the GFP-trap reagent prior to NuPAGE analysis, staining by colloidal coomassie and analysis of excised bands by mass spectrometry. **F)** Acetylated peptides detected by mass spectrometry analysis. Also see Supplemental Figure S2.

### Regulation of the synthetic substrate by sirtuins

We predicted that our synthetic substrate would be regulated by enzymes used to derive the consensus sequence, namely sirtuin HDACs and the Gcn5 HAT. To test whether our substrate was regulated by sirtuin enzymes, we measured the acetylation on our synthetic substrate with 3 consensus repeats (3X) following its purification from yeast strains treated with the sirtuin inhibitor nicotinamide. The acetylation observed on the 3X substrate increased dramatically with nicotinamide treatment and this effect was concentration dependent **(Fig. 2C)**. In contrast, nicotinamide had no impact on acetylation of GFP alone **(Fig. 2C)**. To determine the sirtuins that contribute to this effect, we analyzed the acetylation of the 3X construct in *hst1Δ*, *hst2Δ*, or *sir2Δ* deletion mutants. We observed an increase in acetylation only in *hst2*Δ mutants **(Fig. 2D)**. We next tested acetylation of the 3X substrate in an *hst1Δ hst2Δ sir2Δ* triple mutant, used previously to generate the consensus sequences investigated in this work. We observed a dramatic increase in acetylation of our substrate in this mutant background beyond that observed in *hst2*Δ strains **(Fig. 2D).** Since no single mutant recapitulated the dramatic effect of the sirtuin triple mutant, we suggest that sirtuins act redundantly to deacetylate the synthetic substrate. This is reminiscent of the cooperative sirtuin-dependent regulation of Ifh1 and Sgf73, which we and others described previously (Cai et al. 2013; Downey et al. 2013; Downey et al. 2015). Hst2 is the only sirtuin localized to the cytoplasm, suggesting that the substrate is deacetylated at least partially in this cellular compartment (Perrod et al. 2001; Wilson et al. 2006). Sir2 likely cooperates with Hst1 to deacetylate the fusion substrate in the nucleus.

### Contribution of individual sites to acetylation of tandem consensus sequences

To confirm that acetylation was occurring on the consensus sequence, we purified our fusion protein and mapped acetylation sites following separation by NuPAGE, trypsin digestion and analysis by Orbitrap mass spectrometry. We observed acetylations on the first and second lysine residues when the substrate was purified from sirtuin mutant cells, confirming acetylation of the target sequence *in vivo* **(Fig. 2E,F and Supplemental Fig. S2B, S2C)**. To test if individual lysine residues were equally important we focused on the 2X substrate. We generated variants of the 2X consensus where the first (R1), second (R2) or both (DM) lysine residues were mutated to arginine, which maintains the charge of a lysine residue but cannot be acetylated **(Supplemental Fig. S2D)**. The mutation of only the first lysine residue (R1) resulted in decreased in acetylation as measured by IP-Western analysis **(Supplemental Fig. S2E)**. In contrast, the mutation of the second lysine residue (R2) had little effect **(Supplemental Fig. S2E)**. As expected, mutation of both lysine residues (DM) prevented acetylation altogether **(Supplemental Fig. S2E)**. Since its conversion to arginine resulted in the greatest loss of signal, it appears that the lysine within the first consensus repeat is normally more heavily acetylated than the lysine in the second repeat. It is possible that acetyltransferases have difficulty in acetylating residues very close to the C-terminus of protein sequences. Decreased acetylation observed for the R1 mutant, but not the double mutant, was rescued by mutation of sirtuin enzymes. **(Supplemental Fig. S2E)**, consistent with our results using the 3X substrate (**Fig. 2D**).

### *In vivo* regulation of the consensus sequence by Gcn5

We next tested the contribution of Gcn5 to the acetylation of the 3X synthetic substrate. Acetylation was eliminated in cells lacking the Gcn5 HAT, confirming dependence on this enzyme *in vivo* **(Fig. 3A)**. The regulation of our synthetic substrate by the opposing activities of the Gcn5 HAT and sirtuin HDACs validates the consensus sequences for these enzymes and suggests that target sequences are an important determinant of acetylation. To our knowledge, this is the first demonstration of a portable HAT consensus sequence that directs acetylation *in vivo.*

**Figure 3:**
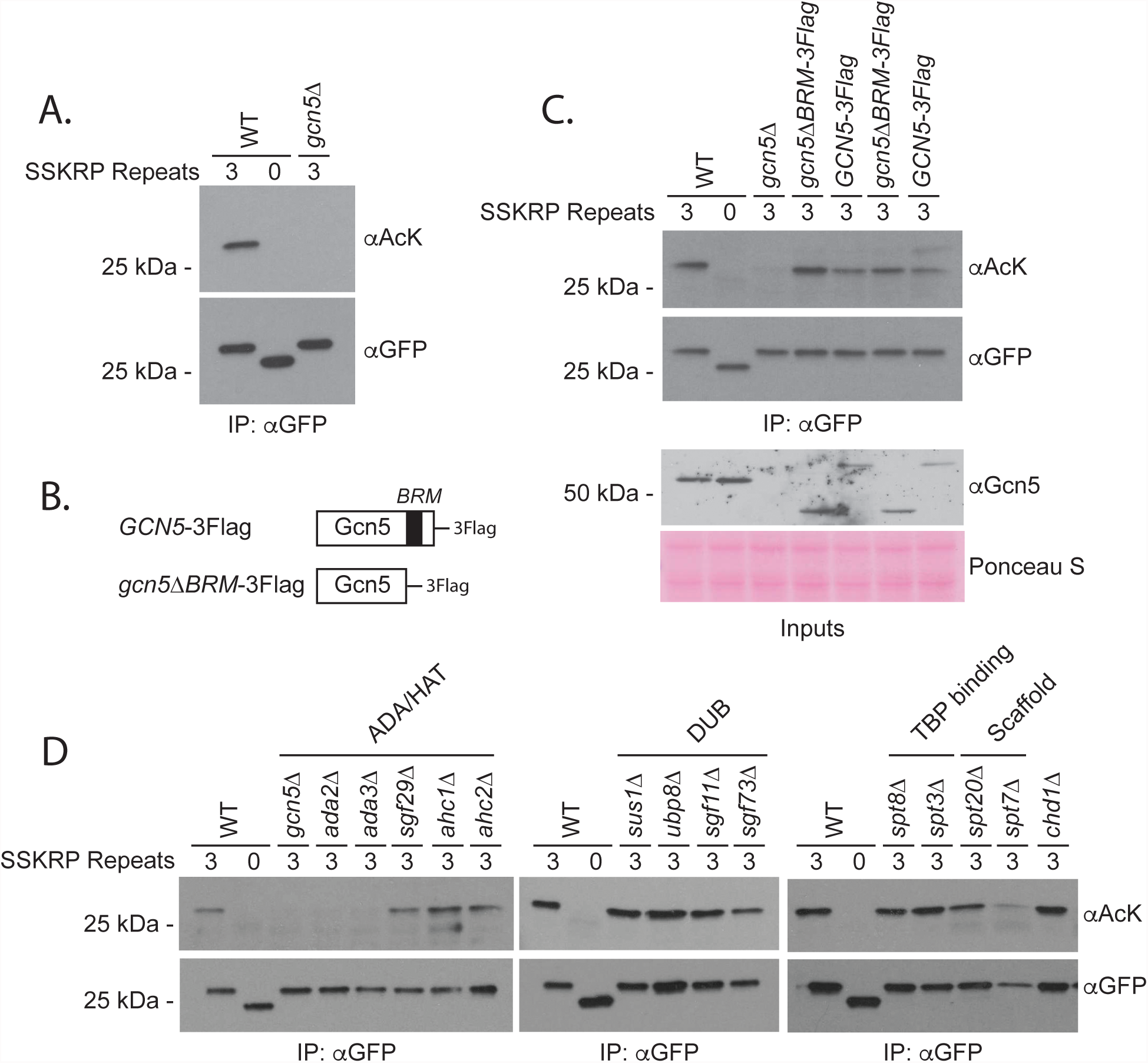
Acetylation of a synthetic substrate is dependent on Gcn5. **A)** The indicated constructs were purified from the strains shown using an antibody against GFP prior to analysis by SDS-PAGE, Western blotting and detection with αAcetyllysine or αGFP. Diluted forms (1/25) of immunoprecipitated protein samples were loaded for GFP detection. **B)** The role of the Gcn5 bromodomain in substrate acetylation was tested using the strains shown. **C)** The indicated constructs were analyzed as in (A) following purification from the strains shown. **D)** The indicated constructs were analyzed in SAGA mutants shown as described in (A).

Previous work suggested that Gcn5’s bromodomain plays a role in regulating acetylation of adjacent lysine residues on histones (Cieniewicz et al. 2014). Whether this is a general property of Gcn5 function that is also applicable to non-histone substrates is unknown. To test this, we assayed the acetylation of the 3X substrate recovered from strains where Gcn5 was mutated for its bromodomain (Gcn5ΔBRM) **(Fig. 3B)**. Unexpectedly, the substrate showed increased acetylation in Gcn5ΔBRM strains, relative to matched controls **(Fig. 3C)**. The increase in acetylation may stem from a moderate increase in Gcn5 expression that was observed in the absence of the bromodomain (**Fig. 3C)**. While these data suggest that the Gcn5 bromodomain does not contribute to the overall acetylation of our construct, we cannot exclude the possibility that it regulates the relative distribution of acetylation marks among individual lysines.

### Regulation of non-histone protein acetylation by SAGA subunits

We next used our synthetic substrate as a tool to test the contribution of individual SAGA subunits to non-histone protein acetylation *in vivo.* Interestingly, expression of the construct varied considerably amongst SAGA mutants **(Supplemental Fig S3A).** Nevertheless, our optimized IP protocol recovered similar levels of protein from SAGA mutants, allowing us to assess the contribution of individual proteins to acetylation of the synthetic substrate. We found that acetylation was largely unaffected by deletion of genes encoding the DUB, TBP-binding and structural proteins, as well as potential SAGA-binding protein Chd1 (Pray-Grant et al. 2005) **(Fig. 3D)**. The lack of effect in *spt7*Δ and *spt20*Δ mutants is particularly intriguing. Although we recovered less substrate from these mutants, the protein that we did recover was acetylated at near wild-type levels **(Fig. 3D)**. The impact of the HAT subcomplex varied depending on the subunit in question. A striking defect in acetylation of our substrate was observed in the absence of HAT submodule proteins Ada2 and Ada3 **(Fig. 3D)**. On the other hand, Sgf29 and ADA-subcomplex specific components Ahc1 and Ahc2 were largely dispensable. We suggest that our synthetic substrate functions as a ‘generic’ target whose acetylation or lack thereof is predictive of intimate effects on Gcn5 or its nearest neighbours. Our substrate could be used to separate modes of regulation that are substrate specific. For example, since Spt7 and Spt20 are required for SAGA stability and transactivator function (Sterner et al. 2002a; Wu and Winston 2002; Lee et al. 2011; Kassem et al. 2017), dependency on these factors could signal that acetylation of non-histone substrates is occurring on chromatin. In support of this idea, Gcn5-mediated acetylation of transcription factor Ifh1, which occurs at promoters of ribosomal protein genes, requires *SPT7* (Downey et al. 2013).

### Ada3 is a global regulator of acetylation

Having identified Ada3 as a potential regulator of non-histone protein acetylation using our synthetic substrate, we carried out acetylome profiling for cells mutated for *ada3*Δ to validate our results test for effects on acetylations proteome-wide **(Fig. 4A)**. We obtained SILAC ratios for 549 acetylated peptides, with 38 showing >2fold downregulation relative to wild-type **(Fig. 4B; Supplemental Table S3)**. GO-term analysis revealed that regulated proteins function predominantly in translation and chromatin-related processes **(Fig. 4C,D)**. These functional categories are reminiscent of what we observed previously for Gcn5 targets (Downey et al. 2015). Included in this group were previously identified Gcn5 targets such as Sgf73 K288 (Downey et al. 2015) and novel targets including Spt16 K464 **(Fig. 4B)**. These data confirm a global role of Ada3 in the regulation of protein acetylation. Perhaps unexpectedly, we also uncovered 88 acetylated peptides that were upregulated >2 fold in *ada3*Δ relative to wild-type cells **(Fig. 4B; Supplemental Table S3)**. GO-term analysis showed enrichment for cytosolic proteins involved in glycolysis and gluconeogenesis **(Fig. 4C,D)**. The functional enrichment was significant when the background used for normalization was limited to proteins recovered in our mass spectrometry experiments **(Supplemental Fig. S3B)**. Moreover, regulated proteins annotated to glycolysis and gluconeogenesis show extensive physical interactions **(Supplemental Fig S3C)**. Upregulated protein acetylations could be the result of indirect effects on other HAT and HDAC enzymes.

**Figure 4:**
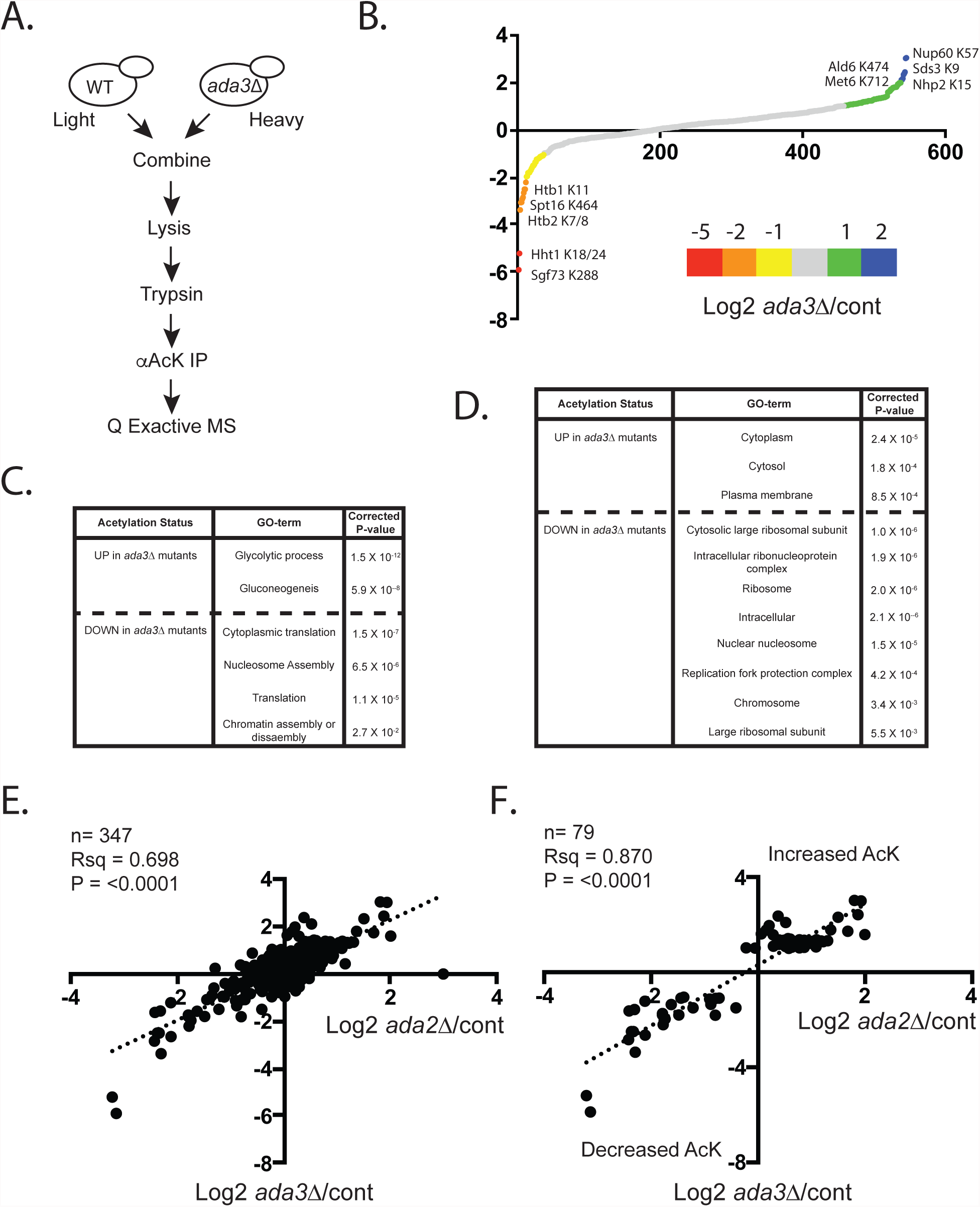
Ada3 regulates acetylations proteome-wide. **A)** Schematic of SILAC-based MS protocol used for acetylome analysis. **B)** Average Log2 fold change for *ada3*Δ/WT for all peptides detected in SILAC experiments. Graph includes combined results of forward and reverse label experiments. **C)** GO-term analysis for ‘Biological Process’ was done using DAVID 6.8. Regulated peptides are >2-fold change in the indicated direction. **D)** GO-term analysis for Cellular Component using DAVID 6.8. Regulated peptides are >2-fold changed in the indicated direction. **E)** Comparison of *ada3*Δ/control versus *ada2*Δ/control ratios for acetylated peptides in Downey *et al* 2015. **F)** As in (E) but just peptides found to be >2fold in *ada3*Δ/control experiments in either direction.

We compared the results of our *ada3*Δ experiments to those of similar experiments performed in *ada2*Δ mutants (Downey et al. 2015) and found significant correlation of the data sets **(Fig. 4E)**. This correlation persisted when unregulated peptides were excluded from the analysis **(Fig. 4F)**. Together the analysis suggests that Ada2 and Ada3 work together to regulate protein acetylation of non-histone substrates by Gcn5.

To investigate the mechanism by which Ada3 impacts protein acetylation, we used a coimmunoprecipitation strategy to compare Gcn5 binding partners in *ADA3* and *ada3*Δ cells. Consistent with our previous findings in *ada2*Δ mutants, Gcn5 failed to bind to SAGA in the absence of *ada3*Δ **(Fig. 5A)**. Our analysis of the synthetic substrate in SAGA mutants suggests that association of the HAT complex with SAGA, lost in the absence of the ADA proteins, is not required for acetylation of non-histone substrates *per se* **(Fig. 3D)**. Yet, as discussed above, SAGA may impact acetylation of some targets. Interestingly, Ada2 is able to retain interaction with Gcn5 in the absence of Ada3, although analysis of input material reveals less Ada2 protein overall **(Fig. 5B).** These data support a model where Ada2 and Ada3 cooperate with Gcn5 to regulate the acetylation of non-histone substrates. In the absence of Ada3, Ada2 can still bind to Gcn5 but this subcomplex is less abundant and incapable of maintaining balanced levels of non-histone protein acetylation **(Fig. 5C)**.

**Figure 5:**
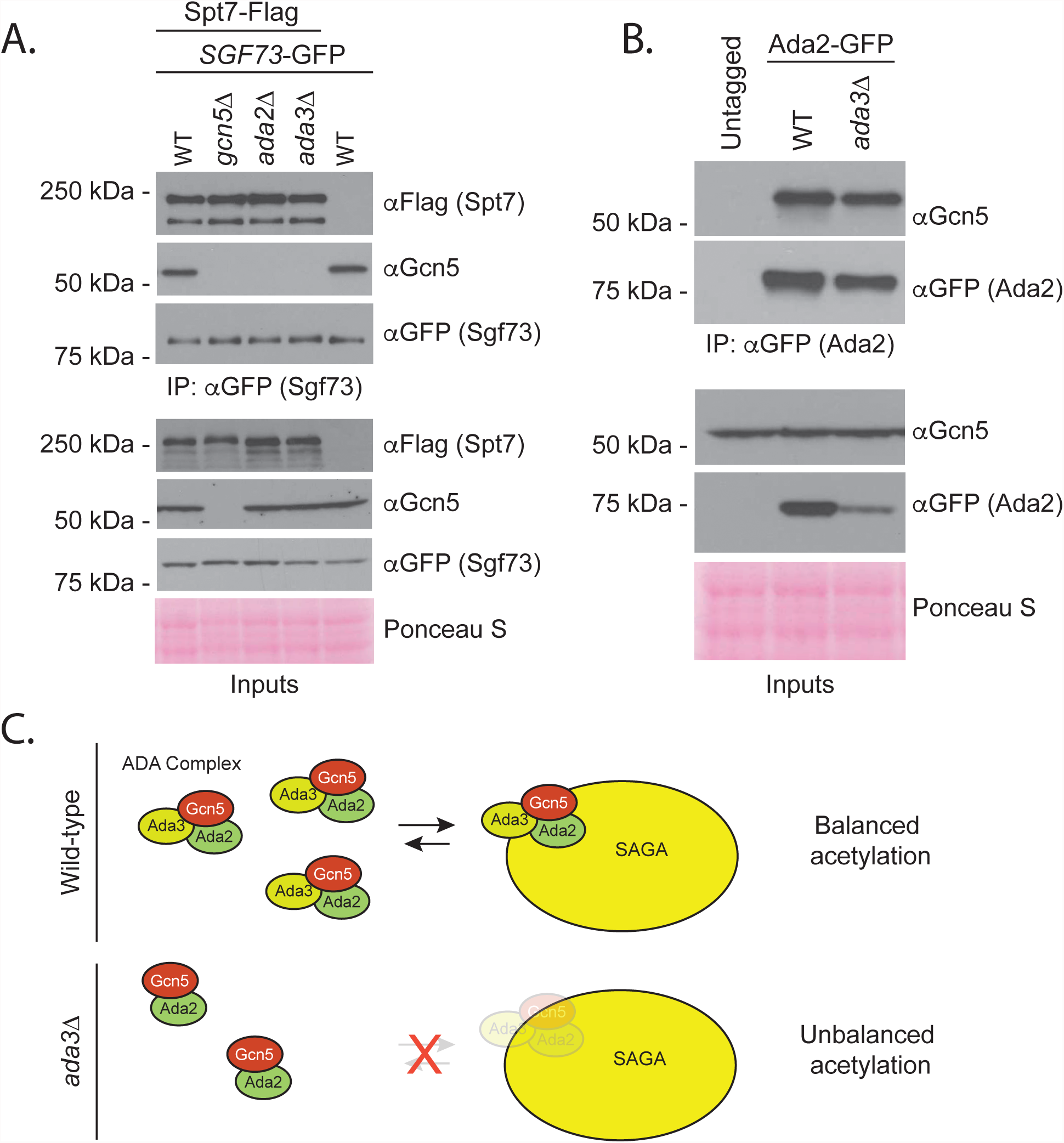
ADA3 mutation prevents Gcn5 binding to SAGA but is permissive for Ada2-Gcn5 interaction. **A)** SAGA subunit Sgf73 was immunoprecipitated via GFP tag in the indicated strains and Spt7-3flag or Gcn5 were detected with the antibodies shown following SDS-PAGE and Western blotting. **B)** Ada2 was immunoprecipitated with a GFP tag in the indicated strains and Gcn5 was detected with an aGcn5 antibody following SDS-PAGE and Western blotting. **C)** Model for Ada3’s role in Gcn5-dependent acetylation of non-histone targets

Altogether our work serves to illustrate the importance of consensus sequences in Gcn5 and sirtuin substrate targeting. Extension of our synthetic biology approach to other HAT and HDAC enzymes across model systems will generate a toolkit to compare and contrast mechanisms of regulation *in vivo.*

## METHODS

### Yeast strains

Yeast strains are in the S288C background and described in Supplementary Table S1. All strains were generated using standard procedures (Longtine et al. 1998; Goldstein and McCusker 1999) and verified using a combination of PCR analysis of colony-purified transformants and Western blotting, where appropriate as described previously (Rossl et al. 2016). Primer sequences used to confirm strains are available upon request. Plasmids, described below, were introduced into yeast using high efficiency lithium acetate transformation followed by selection on synthetic complete (SC) media lacking uracil.

### Plasmids

To construct the entry vector for GFP-consensus constructs, the multi-cloning site from pRS316 was cloned into pRS406-*ADH1*/*CYC1* (A gift from Nicolas Buchler and Fred Cross, Addgene plasmid number 15974) using the restriction enzymes *Kpn*1 (NEB R3142) and *Sac*I (NEB R3156S). GFP or GFP-consensus constructs were generated by amplifying GFP from plasmid pFA6-GFP-His3MX using PAGE-purified oligonucleotides (Eurofins, sequences available upon request). The 5’ oligo included a *Hind*III restriction site. The 3’ oligo included consensus sequences followed by an *Eco*RI site. Constructs and vectors were digested with *Hind*III (NEB R0104S) and *Eco*RI (NEB R0101S) for 90 minutes 37 °C. Agarose gel-purified fragments were ligated using T4 DNA ligase (NEB MO202) prior to transformation into chemically competent DH5α cells (ThermoFisher 18263012) and recovery via plasmid miniprep (Biobasics BS614). Construct sequences were verified via Sanger sequencing (McGill University and Génome Québec Innovation Centre) using primers within the *ADH1* promoter, GFP coding sequence and the *CYC1* terminator. For *SPT2*-GFP plasmids, pRS316 was first digested using *Hind*III-HF (NEB, R3104S) and *Eco*RI-HF (NEB, R3101T) for 15 min at 37 °C. *SPT2*-GFP cassette was amplified using primers providing homology with pRS316. Agarose gel-purified fragments were combined in Gibson assembly mix (NEB, E5510S) and incubated for 15 minutes at 50 °C. Product was then transformed into chemically competent cells (NEB, E5510S) and recovered via plasmid miniprep, followed by Sanger sequencing verification. *spt2-*K166R-GFP mutant plasmid was created by amplifying sequencing primers with overlapping primers to introduce two separate nucleotide changes that add an *Xba*I restriction site (non-coding) in addition to a mutation conferring the desired lysine to arginine change. Reaction mixture was digested with *Dpn*I, transformed into chemically competent cells and then recovered via plasmid miniprep. Plasmids were first confirmed by digestion using *Xba*I (NEB R0145), followed by verification by Sanger sequencing. Plasmids will be made available through Addgene (www.addgene.org).

### Whole cell extract generation and Immunoprecipitation

40-80 OD_600_ equivalents of log phase cells were collected and lysed using acid-washed glass beads in 750 µL chilled IP lysis buffer (50 mM Tris-HCl pH8.0, 5 mM EDTA, 150 mM NaCl, 0.5% NP40) with inhibitors (10 mM Glycerol-2-phosphate, 5 mM NaF (Sigma 201154), 1 mM DTT (BioBasic DB0058), 1.75mM PMSF (Sigma P7626), complete protease inhibitor tablet (without EDTA; Roche 4693132001), 10 mM sodium butyrate (Sigma 303410) and 10 mM NAM (Sigma N3376)). Lysis was carried out in screw cap tubes with 8 timed pulses of 1.5 minutes on a BioSpec Mini Beadbeater with incubation on ice in between bursts. Tubes were punctured with an 18-gauge needle and supernatant was collected in 75 mm tubes (Falcon ref. 352054) via centrifugation, transferred to microfuge tubes and spun 4 minutes at 17000 g. Supernatants were transferred and spun again for 4 minutes at 17000 g before transferring again to a clean microfuge tube. 20-50 µL of cell extract was saved for inputs. Remaining supernatants were incubated at 4°C for 2 hours with 0.5 µL anti-GFP antibody (Abcam ab290) and then another hour with 20 µL washed magnetic beads coupled to Protein A (Bio-Rad 161-4013). Beads, antibody, and bound proteins were recovered on the magnetic Dynarack and washed 3 times in IP lysis buffer, followed by elution in 1-2X SDS sample buffer (with DTT at a final concentration of 100 mM) at 65 °C for 10 minutes. Eluates were transferred to new tubes prior to heating at 100 °C for 5 minutes and analysis via SDS-PAGE.

### Immunoblotting

Unless indicated otherwise, gels were 10% SDS-PAGE with 37.5:1 acrylamide:bisacrylamide (BioRad 1610158). Gels were transferred to PVDF membrane (Biorad 162-0177) at 75 V. All membranes were blocked in 5% milk or BSA in TBS with Tween (0.1%). Primary antibody mixtures were made at a 1:2000 dilution unless otherwise mentioned, in either 5% milk or BSA in TBS-T with 0.01 % sodium azide. Membranes were incubated at 4 °C overnight, washed 3 × 10 minutes with TBS-T before probing with HRP-coupled secondary antibodies (also made in 5% milk or BSA, at 1:10,000 dilution) for 30-50 minutes. Blots were then washed an additional 3 X in TBS-T for 10 minutes each prior to application of ECL reagent (Millipore) and exposure to autoradiography film (Progene). Product numbers and concentrations of antibodies used are summarized in Supplementary Table S2.

### Mass spectrometry

#### For determination of acetylation sites on GFP-consensus constructs

Indicated constructs were immunoprecipitated via GFP Trap (Chromotek). Bound proteins were eluted with SDS-PAGE sample buffer and analyzed on a NuPAGE Novex Bis-Tris (4-12%) gel (ThermoFisher NP0336BOX) run at 200V for 50 minutes according to manufacturer’s instructions and as described elsewhere (Bentley-DeSousa et al. 2018). Staining was with Invitrogen Colloidal blue staining kit (LC6025) following manufacturer’s directions for Novex Bis-Tris gels. Preparation of excised gel slices were carried out as described (Shevchenko et al. 2006).

ELITE-LC MS/MS was completed as previously described with minor modifications (Starr et al. 2017). Briefly, our analysis employed an Ekspert NanLC 400 (Eksigent, Dublin, CA) and an Orbitrap ELITE MS (ThermoFisher Scientific, San Jose, CA, USA). The MS was operated in the positive ion mode. Peptides were resuspended in 30 µL of 0.5 % formic acid prior to injection into an analytical column of 75 µM internal diameter and packed with 1.9 µM C18 Resin (Dr. Maisch, CmbH, Ammerbuch, Germany). Elution was with a flow rate of 300 nL/min. A 120 minute gradient of 5-30 % (v/v) acetonitrile with 0.1 % (v/v) formic acid was used. The heating capillary was set at 300 °C. The spray voltage was fixed at 2.2 kV. The MS scan used ranged from 350-1750 m/z. The MS/MS scan was conducted on the 20 most intense ions. Exclusion duration was 90 seconds with one repeat count and a 30 second repeat duration.

#### For acetylome analysis

SILAC labeling for paired WT and *ada3*Δ mutant cells, cell lysis, chemical treatments, trypsin digestion (ThermoFisher 90058), anti-AcK IP (ImmuneChem ICP0380), elution, and peptide purification prior to mass-spectrometry analysis were as previously described (Downey et al. 2015). High-performance liquid chromatography electrospray ionization tandem mass spectrometry (HPLC-ESI-MS/MS) for yeast acetylome analyses was completed using the Q Exactive mass spectrometer (ThermoFisher Scientific, San Jose, CA). Conditions used were similar to those described elsewhere (Zhang et al. 2018). Briefly, the Q Exactive instrument was operated in positive ion mode. Peptides immunoprecipitated with anti-acetyllysine antibody were first resuspended in 0.5% (v/v) formic acid and injected onto a 75µM internal diameter analytical column packed with 1.9µm C18 resin (Dr. Maisch, GmbH, Ammerbuch, Germany). Peptides were eluted using a 200nL/min flow rate. A 120 minute gradient was used with increasing acetonitrile concentration (5-30% (v/v), 0.1% (v/v) formic acid). The MS scan employed ranged from 300 to 1800 m/z with subsequent selection of the 12 most intense ions for data-dependent MS/MS scan. A dynamic exclusion repeat count of 2 and repeat exclusion duration of 30 seconds were used.

### Database searches

Xcalibur software (ThermoFisher Scientific, San Jose, CA) was used to acquire data. Following acquisition, a search was performed using MaxQuant software version 1.5.3.30 (Cox et al. 2009) against a *Saccharomyces cerevisiae* database (downloaded from Uniprot 2017/02/09). Parameters used were: multiplicity of two (heavy label: Lys8); trypsin digest, a maximum of two missed cleavages, fixed modification of cysteine carbamidomethylation; variable modifications of methionine oxidation, acetyllysine and N-terminal acetylation; minimum peptide length of seven amino acids; 0.5 Da for ion mass tolerance; peptide and protein false discovery rate fixed at 1%. For in-gel analysis, the GFP-3X consensus fusion sequence was added to the database.

### Bioinformatics analyses

GO-term enrichments were determined using DAVID version 6.8 (david.ncifcrf.gov) with *S. cerevisiae* as the background and with default settings (Huang et al. 2007; Huang da et al. 2009). Network diagrams were constructed using Genemania (Genemania.org) (Warde-Farley et al. 2010).

### Antibody generation

Antibody protocols were developed with the intention to generate a reagent that recognizes the critical features of the acetylated consensus without being specific to an exact amino acid sequence. Hybridoma clone A1504705 was developed by immunizing four BALB/c mice with 25 µg of KLH conjugated peptide AAASAK(ac)RPAAA prepared in Complete Freund’s adjuvant (Sigma-Aldrich, Cat No: F5881). Two weeks later, each mouse was boosted with 12.5 µg of KLH conjugated peptide GAPANK(ac) RPRRG prepared in Incomplete Freund’s adjuvant (Sigma-Aldrich, Cat No: F5506), followed by another boost with 12.5 µg of KLH conjugated peptide SSVSYK(ac)RVCGG prepared in Incomplete Freund’s adjuvant two weeks apart. Three days after the boosting, sera of immunized mice were collected and tested against BSA conjugated peptides AAASAK(ac)RPAAA, GAPANK(ac)RPRRG, SSVSYK(ac)RVCGG, and AAASAKRPAAA in ELISA. Mouse with best response to first three peptides was subsequently boosted with 10 ug of each KLH conjugated peptides AAASAK(ac)RPAAA, GAPANK(ac)RPRRG, and SSVSYK(ac)RVCGG. Lymphocytes from the mouse received final antigen boost were harvested three days later and fused with myeloma cells sp2/0 using GenomOne Kit (Cosmo Bio, Cat No: ISK-CF-001-EX). Clone A1504705 was selected based on its reactivity to BSA conjugated peptides GAPANK(ac)RPRRG, SSVSYK(ac)RVCGG, and AAASAK(ac)RPAAA, but not to BSA conjugated peptide AAASAKRPAAA in ELISA.

Hybridoma cell culture supernatants were collected and the antibody was purified via a Protein G column (Sigma-Aldrich, Cat No: GE17-0618-01). After supernatant binding, the resin was washed with PBS (pH 7.2) and antibodies were eluted with 50 mM of diethanolamine (pH = 11.0) (Sigma-Aldrich, Cat No: 31589). Subsequently, eluted antibodies were neutralized by 1M Tris (pH = 8.0). The antibody was prepared by dialyzing against PBS (pH = 7.2) with 0.09% of Sodium Azide (Sigma-Aldrich, Cat No: 71289).

## ACKNOWLEDGEMENTS

We thank members of the Downey lab for critical reading of the manuscript. We thank members of the Figeys Lab for assistance with protein purification and mass spectrometry. This project is supported by NSERC grant RGPIN-2016-05015 to MD.

## FIGURES & LEGENDS

**Supplemental Figure 1:**
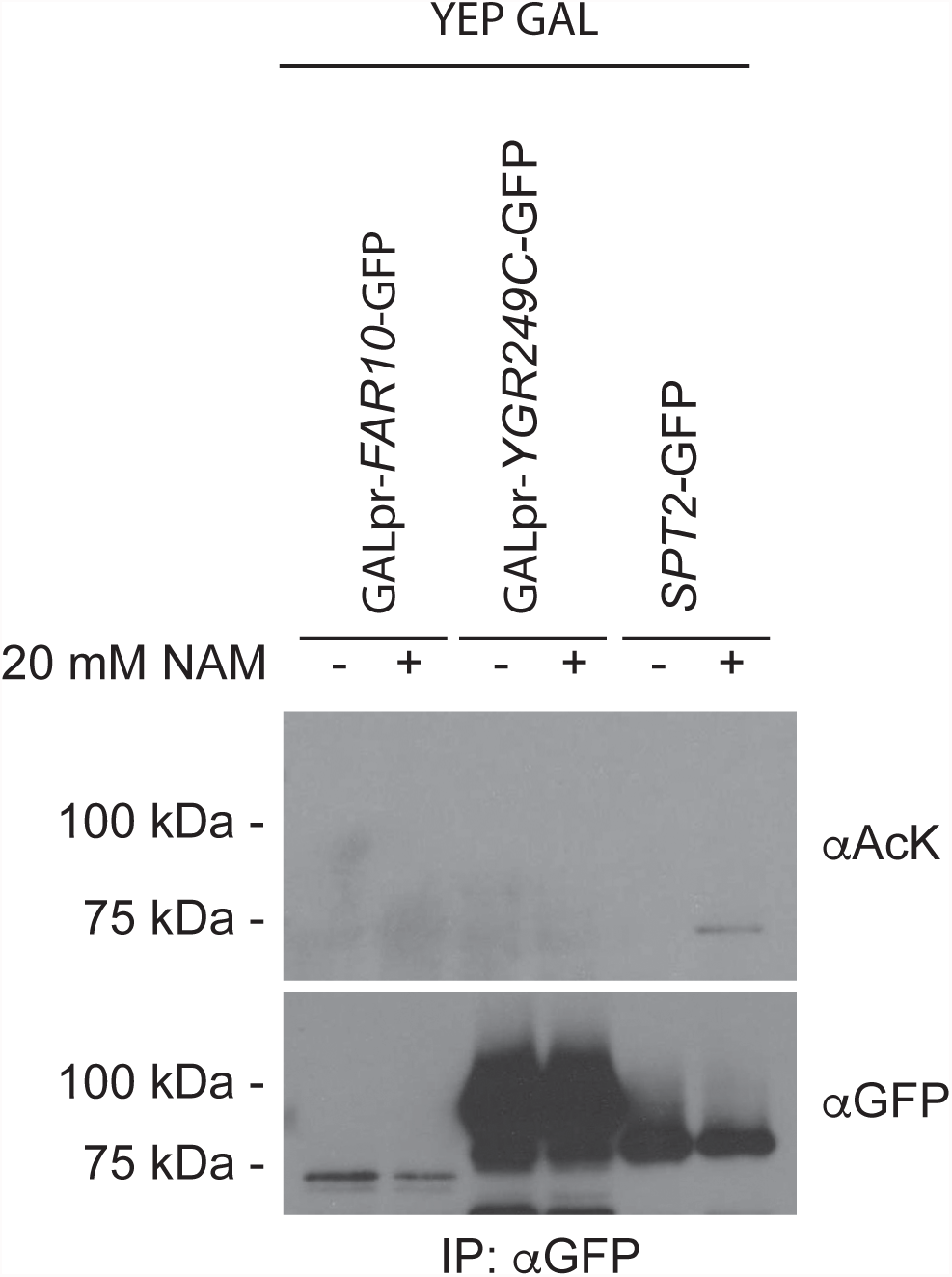
Far10-GFP and Ygr249c are not acetylated. The indicated fusion proteins immunoprecipitated from the strains shown after GAL induction. Eluates were analyzed with the indicated antibodies following SDS-PAGE and Western blotting.

**Supplemental Figure 2:**
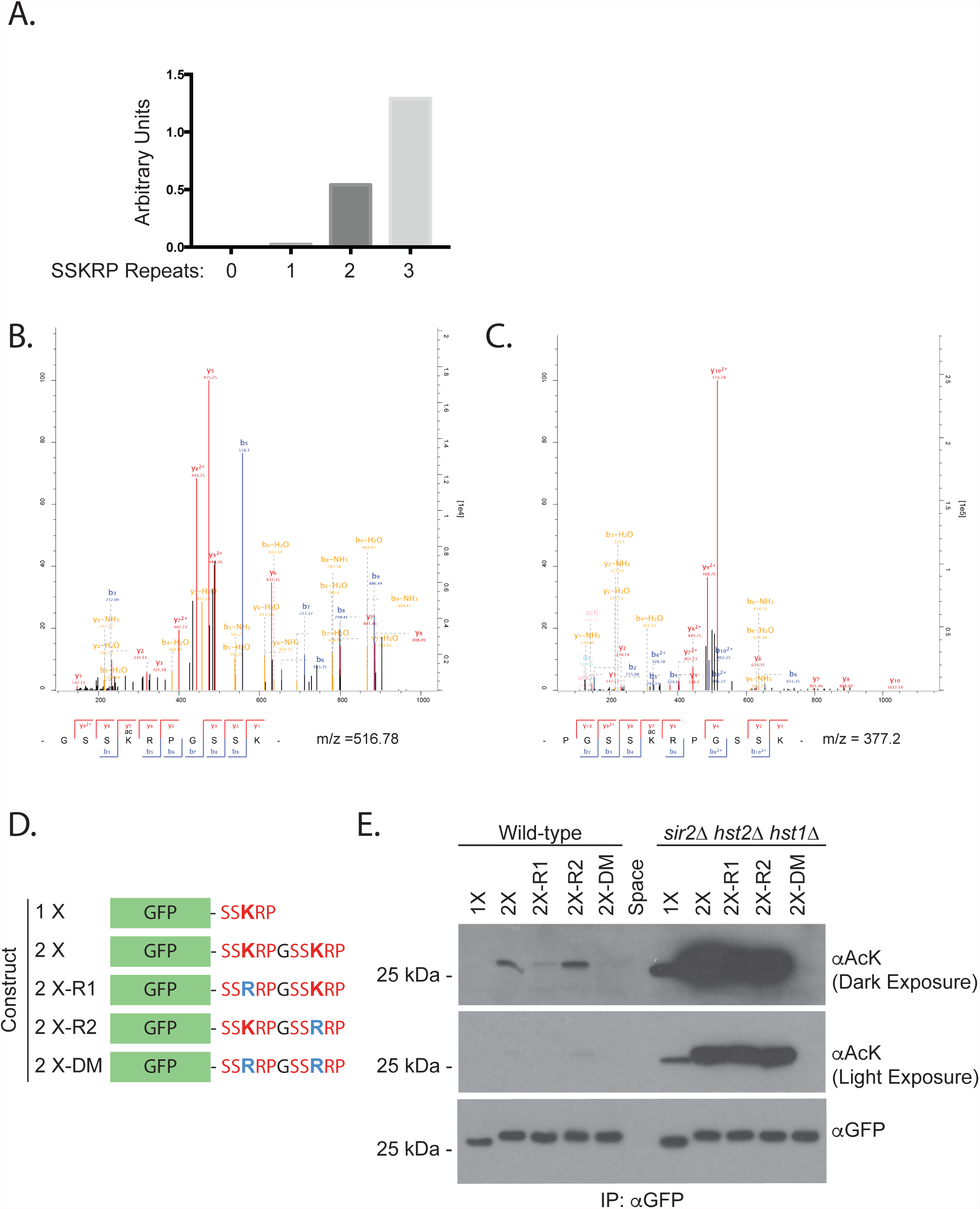
Acetylation of a synthetic substrate *in vivo*. **A)** A lighter exposure of the blot shown in Figure 2B was quantified using ImageJ. Acetylation was normalized to the level of GFP protein and all values normalized to strain expressing GFP alone via subtraction of the background signal in this lane. **B)** and **C)** MS/MS spectra for the indicated acetylated peptides. **D)** 2X fusion constructs used to test the contribution of individual lysine residues to signal observed with monoclonal aAcetyllysine antibodies. **E)** The constructs shown in (D) were immunoprecipitated with an antibody against GFP and analyzed with the indicated antibodies following SDS-PAGE and Western blotting. Diluted forms (1/25) of immunoprecipitated protein samples were loaded for GFP detection.

**Supplemental Figure 3:**
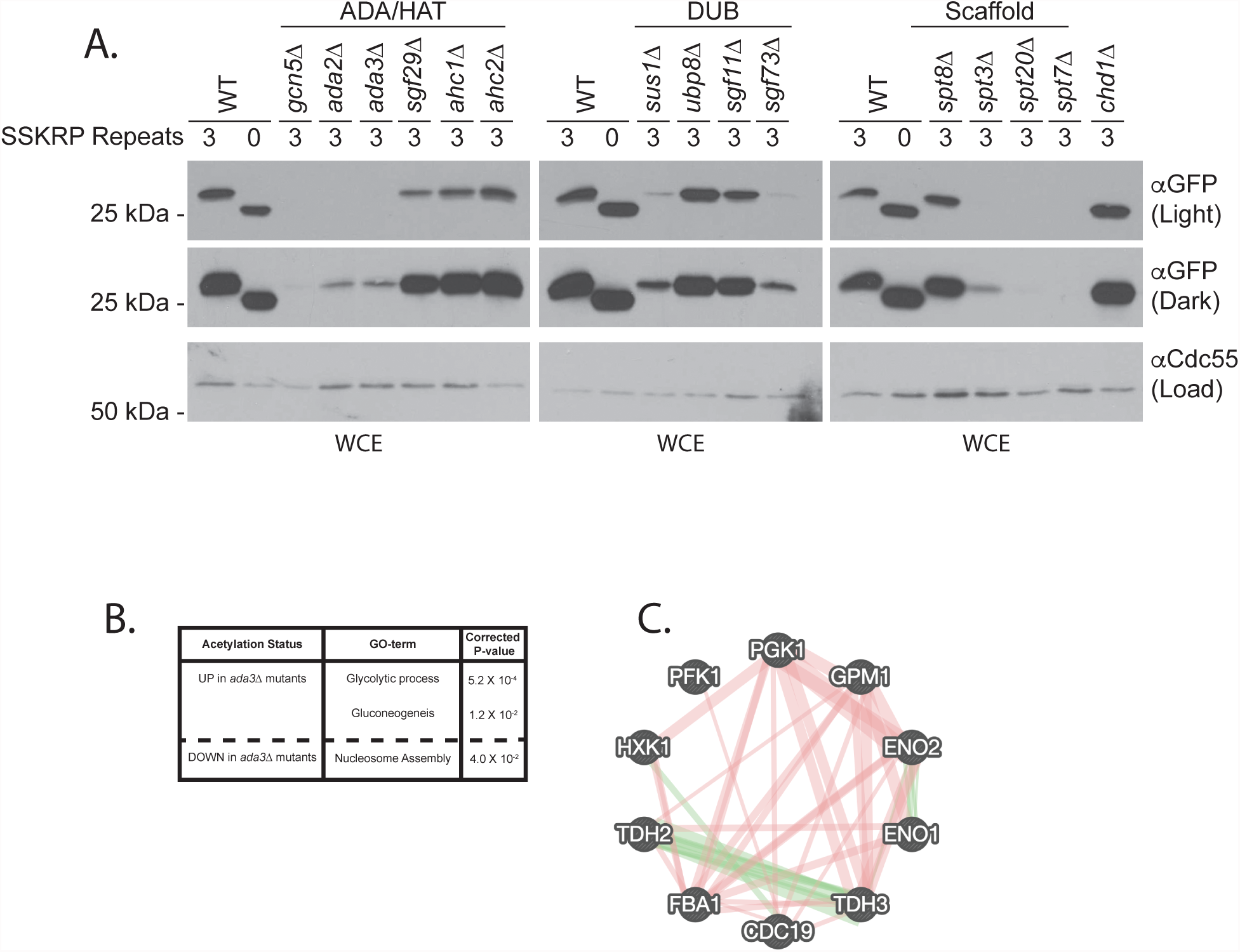
**A)** Input analysis for 3X substrate expression in SAGA mutants. An antibody against Cdc55 was used as a loading control. **B)** Biological process GO-term analysis normalized for recovery to acetylated proteins. **C)** Genetic (green) and physical interactions (red) between glycolytic proteins with acetylations upregulated in *ada3*Δ cells.

## Excel files

**Supplemental Table S1:** Yeast strains used in this study

**Supplemental Table S2:** Antibodies used in this study

**Supplemental Table S3:** SILAC ratios for acetylome profiling of *ada3*Δ cells

